# Improving the accuracy of pose prediction by incorporating symmetry-related molecules

**DOI:** 10.1101/2024.09.21.614298

**Authors:** Sree Hima, Chandran Remya, K.V. Dileep

## Abstract

Accurate prediction of biologically relevant binding poses is crucial for the success of computer-aided drug development. In this study, we describe a general strategy to enhance the precision of pose prediction in molecular docking by incorporating symmetry-related molecules (SRMs). Our objective was to demonstrate the significant impact of SRMs on the accuracy of pose prediction. To achieve this, we evaluated our method on high-quality protein-ligand complex structures, focusing on the presence and absence of SRMs during molecular docking studies. We have extracted the co-crystal ligands from the selected crystal structure and were redocked in presence and absence of SRM to assess their influence. Additionally, we calculated the free energy of the docked poses using the Molecular Mechanics Generalized Born Surface Area (MM-GBSA) method, comparing the results in the presence and absence of SRMs. The findings revealed that redocking performed in the presence of SRMs significantly improved the prediction of biologically significant/crystallographically relevant poses. Consequently, our proposed strategy offers a robust method for enhancing pose prediction in current molecular docking programs, potentially leading to more effective and reliable drug development processes.

## INTRODUCTION

Many drug development campaigns heavily rely on a variety of computational approaches, such as high-throughput virtual screening and molecular docking, to forecast the interactions between ligands and receptors[1-4]. Often the protein-ligand conformations and various sampling algorithms used in the docking programs contributes to the accurate prediction of biologically relevant binding poses [4, 5]. Some of the widely used molecular docking programs are GOLD[6], Glide[7], FlexX[8], AutoDock[9], AutoDock Vina[10], PLANTS[11], rDock[12].Each of these programs employs different algorithms and scoring functions to enhance the accuracy and reliability of their predictions. By leveraging these advanced computational tools, drug developers can improve the efficiency of the drug discovery process, identifying promising candidates more quickly and effectively.

Despite substantial advancements, predicting biologically relevant /crystallographically significant poses remain a challenging endeavour due to several factors such as: **1**. Hurdles in identifying high-quality receptor structures for molecular docking, **2**. Inadequate sampling and scoring algorithms, **3**. Insufficient consideration of solvation effects. The choice of receptor structures, especially incorporating multiple receptor conformations (MRCs), can enhance the accuracy of the pose prediction and virtual screening performances[13]. To improve scoring methods, factors such as shape complementarity, solvation effects, and the entropy contributions of both receptors, ligands and solvents should be considered[14, 15]. Additionally, pose prediction can be significantly improved by integrating structure-based pharmacophores and interaction fingerprints [16], ligand 3D shape similarity [17], and hybrid docking approaches [18]. Also by incorporating highly ordered water molecules, instead of using distributed solvation (i.e., dielectrics), can also enhance pose prediction accuracy [19].

To address the challenges in pose prediction and to generate biologically relevant or crystallographically significant poses, we hypothesized that incorporating symmetry-related molecules (SRMs) from the crystal packing during receptor preparation could enhance the accuracy of pose predictions. A SRM refers to a molecule that is related to another by a symmetry operation in a crystal structure. To validate our hypothesis, we studied the role of SRMs in molecular docking, free energy calculations, and molecular dynamics simulations.

## MATERIALS AND METHODS

### Selection of PDB entries

To study the influence of SRMs on pose prediction, we used a dataset derived from the Protein Data Bank [20], comprising 300 structures with resolutions ranging from 1.00 to 3.50 Å. We previously used this dataset to investigate ambiguous electron density around crystal ligands. Through visual inspection of all collected structures, we selected ligands protruding towards the solvents and applied solvent-accessible surface area (SASA) as filtering criteria. Using PyMOL, we generated symmetry mates to determine whether SRMs directly interact with the crystallographic ligand or not.

### Calculation of SASA

We calculated the SASA of crystallographic ligands to identify those extending towards the solvent. To specifically determine the SASA of these ligands (especially the ligand portion that is protruded towards the solvent), we used the following formula and selected PDB entries with a ligand SASA of ≥ 15% for further analysis.

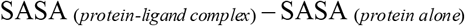

The SASA of the protein-ligand complex was calculated using residue analysis in Schrödinger Maestro, while the SASA of the ligand alone was determined using QikProp.

### Generation of symmetry related molecules

To identify the ligand contacts with SRM, we used COOT [21]and PyMOL [22]. First, we generated symmetry mates with a 4 Å cut-off to understand the orientation of SRM in the crystal packing. We then removed ligand molecules from the symmetry mates to prevent interference with the calculations. PDB entries were selected for further studies if the SRM directly interacted with the crystallographic ligand.

### Protein and ligand preparations for docking studies

The candidate PDB entries in which the ligands have a direct interaction with SRM were prepared with and without SRMs using the protein preparation wizard of Schrödinger Maestro. As a part of preparations, we removed all water molecules from the crystal structure. During preparation, hydrogens were added to polar atoms, partial charges were assigned to metal ions, bond orders were corrected, disulfide bonds were assigned, and energy minimization was performed using the OPLS-2005 force field. The minimization process was terminated when the root mean square deviation (RMSD) of the minimized structure exceeded 0.30 Å compared to the crystal structure. Ligand molecules in the respective crystal structures were prepared using the LigPrep module of Schrödinger Maestro, generating multiple conformations of the ligands at pH 7 ± 2. We then performed molecular docking, with ligands docked to their respective targets in the presence and absence of SRM. Extra Precision (XP) was used for docking studies, and poses with the best docking scores were retained for further analysis.

### Free energy estimation of the ligands in the presence and absence of SRMs

Using the Prime MM-GBSA in Schrödinger Maestro, the free energy of binding was calculated for the docked poses. The protein-ligand complexes both in the presence and absence of SRM were taken for the calculation.

### Molecular dynamic simulation

We also performed molecular dynamics (MD) simulations to evaluate the stability of the docked pose in the presence and absence of SRMs. All MD simulations were performed by Desmond 2021 (GPU version) using the OPLS force field. Prior to MD simulations, the system was set up using the “System Builder” in Desmond. The protein-ligand complex was immersed into an orthorhombic box with dimensions of 10 × 10 × 10 Å containing TIP3P water molecules. A respective number of ions are added to neutralize the system. The MD simulation was performed at a constant temperature of 300 K and the duration of the simulation was 100ns.

## RESULT AND DISCUSSION

### Systematic evaluation of the collected structures

In this study, we examined an in-house dataset consisting of 300 structures with resolutions ranging from 1.05 to 3.50 Å. Although these structures were originally used to study ambiguous electron densities around crystal ligands, we repurposed them for our current investigation. We systematically sorted these 300 structures using various approaches to facilitate our analysis. Initially, our visual inspection allowed us to shortlist approximately

100 structures where the ligands protruded towards the solvent. To further validate the solvent exposure of these ligands, we calculated their SASA. We selected ligands with a SASA of ≥ 15%, which resulted in the elimination of 70 candidate PDB entries.

Next, we generated the SRM and evaluated its interaction with the respective crystallographic ligands. This step led to the exclusion of 19 additional structures that lacked interactions with the SRM. Ultimately, we were left with a dataset of 11 structures, all of which had resolutions below 2.2 Å. At this resolution, we expected the electron density around the ligands to be sufficiently clear.

To ensure the accuracy of ligand fitting, we also evaluated the 2Fo-Fc electron density maps of the selected ligands. The details of the structures and the SASA values of the ligands in the selected PDB entries are shown in **Tables 1 and 2**, respectively. The entire workflow is illustrated in **Figure 1**.

**Table 1:** Details of PDB Structures used in the current study.

| Sl No | PDB ID | Name of structures | Resolution in Å | Space group | $R_{work}$ | $R_{free}$ |
| --- | --- | --- | --- | --- | --- | --- |
| 1. | 5EI4 | First domain of human bromodomain BRD4 in complex with inhibitor 8-(5-Amino-1H-[1,2,4]triazol-3-ylsulfanylmethyl)-3-(4-chlorobenzyl)-7-ethyl-3,7-dihydropurine-2,6-dione | 1.05 | P 2<sub>1</sub> 2<sub>1</sub> 2<sub>1</sub> | 0.171 | 0.192 |
| 2. | 3VNG | Crystal Structure of Keap1 in Complex with Synthetic Small Molecular based on a co-crystallization | 2.10 | P 2<sub>1</sub> 2<sub>1</sub> 2<sub>1</sub> | 0.196 | 0.225 |
| 3. | 2I4H | Structural studies of protein tyrosine phosphatase beta catalytic domain co-crystallized with a sulfamic acid inhibitor | 2.15 | C 1 2 1 | 0.169 | 0.246 |
| 4. | 3KRR | Crystal Structure of JAK2 complexed with a potent quinoxaline ATP site inhibitor | 1.80 | C 2 2 2<sub>1</sub> | 0.167 | 0.206 |
| 5. | 3PBB | Crystal structure of human secretory glutaminyl cyclase in complex with PBD150 | 1.95 | H 3 | 0.181 | 0.239 |
| 6. | 7F7W | JAK2-JH2 (JAK2/FLT3 inhibitor) | 1.83 | P 1 2<sub>1</sub> 1 | 0.207 | 0.238 |
| 7. | 1Q0Z | Crystal structure of aclacinomycin methylesterase (RdmC) with bound product analogue, 10-decarboxymethylaclacinomycin A(DcmA) | 1.95 | P 1 2<sub>1</sub> 1 | 0.157 | 0.185 |
| 8. | 6GL8 | Crystal structure of Bcl-2 in complex with the novel orally active inhibitor S55746 | 1.40 | P 2<sub>1</sub> 2<sub>1</sub> 2<sub>1</sub> | 0.178 | 0.194 |
| 9. | 1MMQ | Matrilysin complexed with Hydroxamate inhibitor | 1.90 | P 3<sub>1</sub> 2 1 | 0.179 | NR* |
| 10. | 5ACL | EGFR kinase domain mutant "TMLR" with compound 24 | 1.49 | P 2<sub>1</sub> 2<sub>1</sub> 2 | 0.158 | 0.187 |
| 11. | 6G0G | KRAS-169 Q61H GPPNHP | 1.48 | P 2<sub>1</sub> 2<sub>1</sub> 2 | 0.193 | 0.220 |

NR\*: Note Reported

**Table 2:** Solvent-accessible surface area of the ligands present in the selected PDB Structures.

| **Sl No** | **Ligand name as per the PDB entry** | **PDB ID** | **Total SASA of ligand in Å<sup>2</sup>** | **% Of SASA in Å<sup>2</sup>** |
| --- | --- | --- | --- | --- |
| 1. | 5NV | 5EI4 | 672.413 | 21 |
| 2. | FUU | 3VNG | 640.158 | 42.34 |
| 3. | UA1 | 2I4H | 825.092 | 42.41 |
| 4. | DQX | 3KRR | 811.189 | 16.02 |
| 5. | PBD | 3PBB | 637.913 | 20.24 |
| 6. | 36H | 7F7W | 832.607 | 18.22 |
| 7. | AKA | 1Q0Z | 1115.222 | 26.30 |
| 8. | F3Q | 6GL8 | 899.223 | 33.59 |
| 9. | RRS | 1MMQ | 750.370 | 33.33 |
| 10. | SAS | 5ACL | 688.272 | 29.70 |
| 11. | SAS | 6G0G | 688.272 | 26.69 |

**Figure 1.**
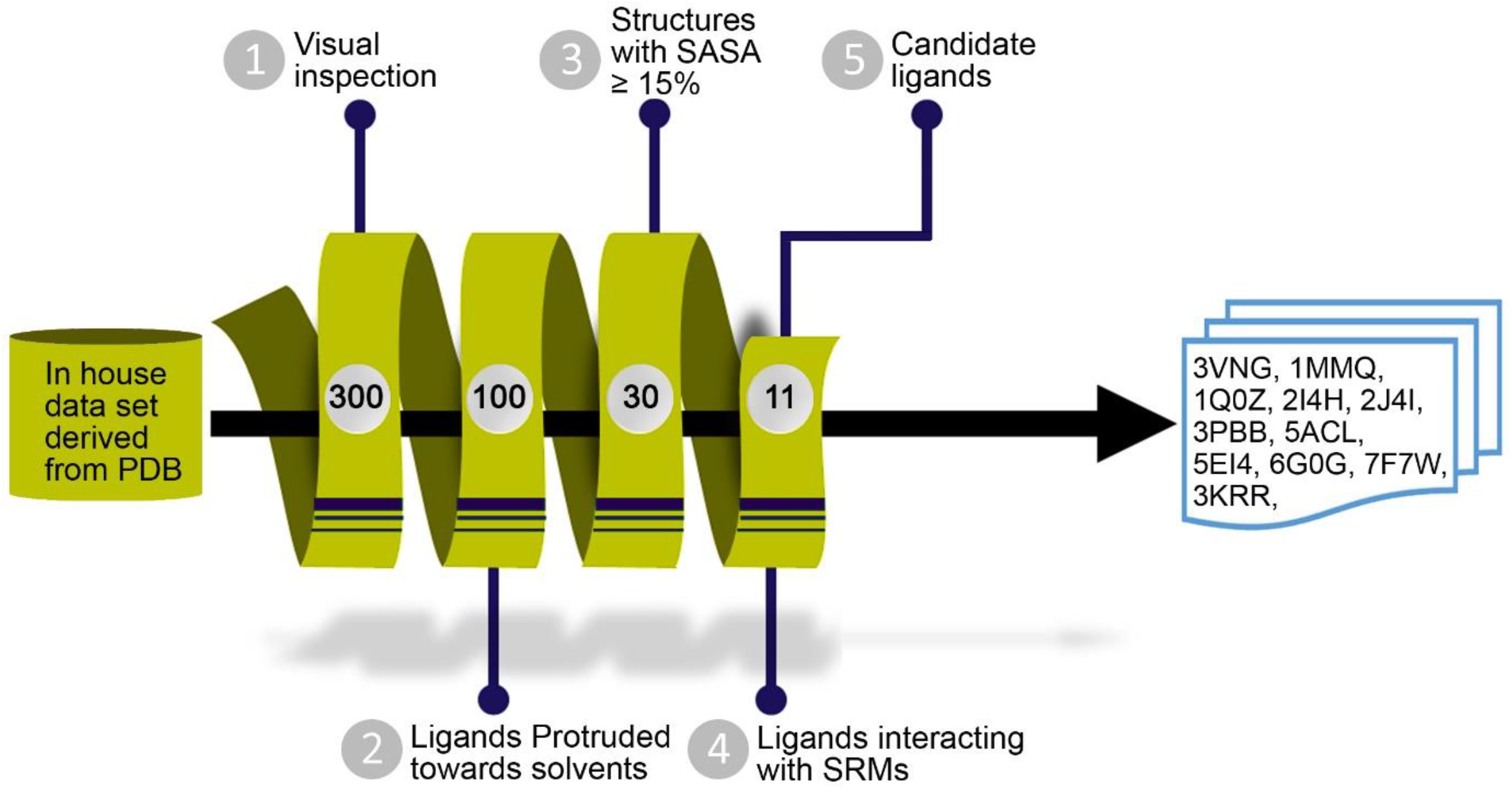
Comprehensive workflow for selecting protein structures for the study. Each step of the process is detailed, including the number of PDB entries shortlisted at each stage. The PDB IDs of the final 11 selected entries are also provided.

### SRM helps to determine the biologically relevant poses

Our molecular docking studies, conducted both in the presence and absence of SRMs, revealed a crucial role for SRMs in retaining biologically relevant/crystallographically significant poses. While comparing the docked poses with crystal poses, significant deviations in the orientation of the ligands were observed for most of the selected structures except 5ACL, 6GL8 and 6G0G in the absence of SRMs **(Figure 2A)**. On the other hand, in the presence of SRMs, the docking programs were able to generate binding poses closely resembling the crystal poses **(Table 3)**.

**Table 3:** The RMSD and MM-GBSA calculated for the ligand in the presence and absence of symmetry-related molecules.

| Sl No | PDB ID | RMSD in Å |  | MM-GBSA |  |
| --- | --- | --- | --- | --- | --- |
|  |  | Symmetry |  | Symmetry |  |
|  |  | - | + | - | + |
| 1. | 5EI4 | 7.12 | 0.84 | -76.19 | -87.63 |
| 2. | 3VNG | 8.26 | 0.89 | -53.42 | -56.01 |
| 3. | 2I4H | 7.93 | 0.29 | -62.88 | -107.24 |
| 4. | 3KRR | 8.66 | 0.49 | -104.96 | -119.52 |
| 5. | 3PBB | 7.59 | 0.25 | -77.77 | -100.89 |
| 6. | 7F7W | 8.54 | 1.00 | -86.03 | -92.82 |
| 7. | 1Q0Z | 8.71 | 0.29 | -137.15 | -141.28 |
| 8. | 6GL8 | 6.71 | 0.43 | -102.21 | -113.36 |
| 9. | 1MMQ | 6.32 | 0.96 | -98.97 | -112.05 |
| 10. | 5ACL | 8.09 | 0.36 | -53.02 | -58.25 |
| 11. | 6G0G | 7.77 | 1.00 | -31.48 | -66.36 |

**Figure 2.**
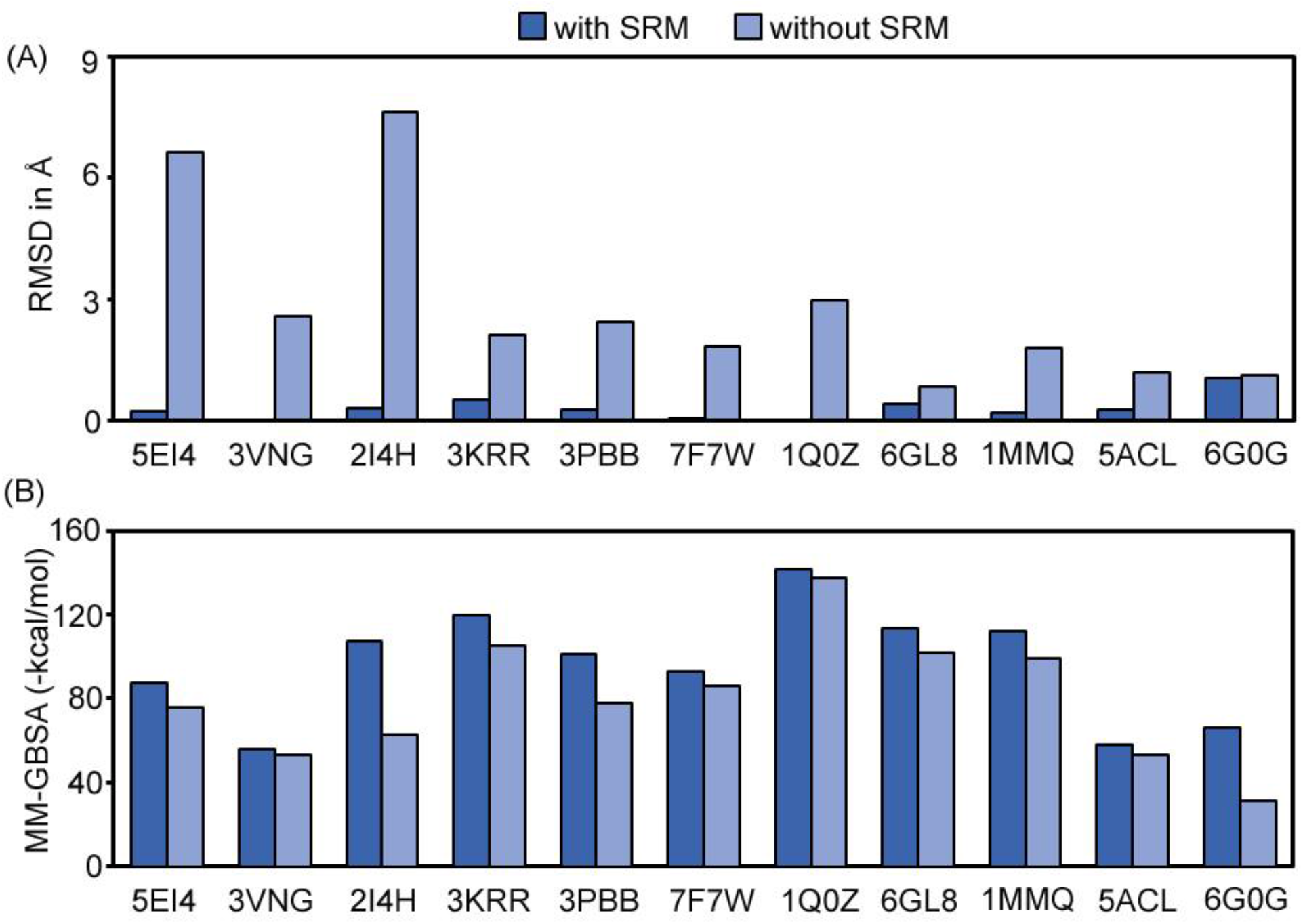
Comparative analysis of ligand position and energy in selected PDB entries with and without SRMs. (A) RMSD of ligands obtained from molecular docking studies. (B) MM-GBSA binding free energy calculations for the selected docked poses. In both figures, the dark blue represents the presence of SRMs, while light blue represents their absence.

Furthermore, an analysis of the binding free energies indicated that all entries exhibited better (more negative) binding free energies in the presence of SRMs when compared to their absence **(Figure 2B)**. This suggests that the binding poses were correctly predicted in the presence of SRMs, thereby favoring the biologically relevant/crystallographically significant conformations. This study underscores the importance of considering SRMs in molecular docking to achieve accurate and reliable predictions of ligand binding poses.

### Mode of action of ligands: crystal structure vs. docked structures in the presence and absence of SRM

To delve deeper into the molecular mechanisms underpinning the ligand interaction with the protein structure, we analyzed the docking results in the absence of SRMs, focusing on the highest RMSD cases. The docking studies revealed multiple interactions for the ligands in the absence of SRMs. In all selected cases, the docked poses of ligands in the presence of SRMs closely resembled the crystal pose **(Figure 3)**.

**Figure 3.**
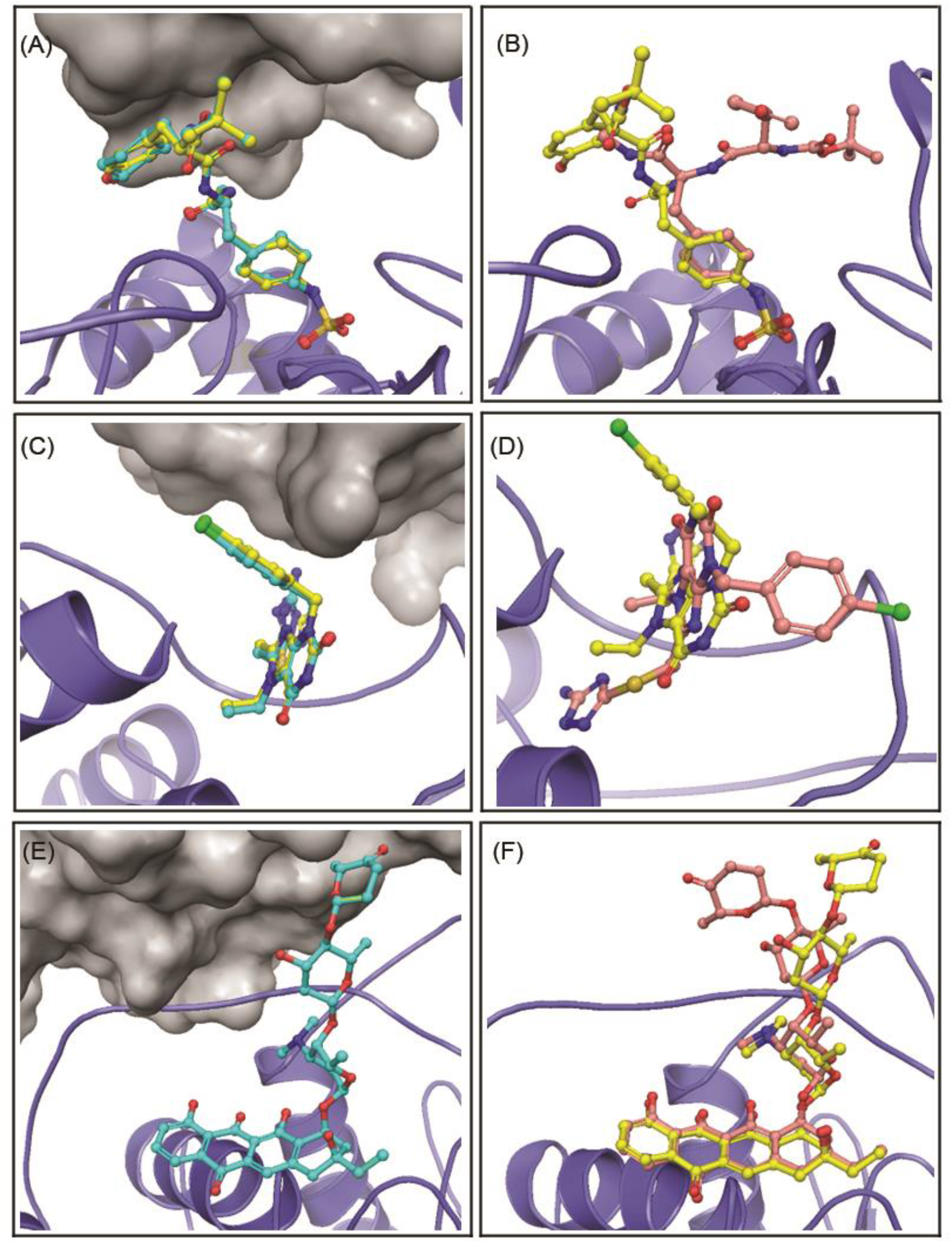
Structural comparison of binding modes of the selected candidates: A, C, and E represent the superposition of docked (in presence of SRM) and crystal poses in 2I4H, 5EI4 and 1QOZ respectively. Whereas B, D and F represents the superposition of docked ligand (in absence of SRM) and crystal pose in 2I4H, 5EI4 and 1QOZ respectively. Proteins receptor is represented in slate blue colour and grey surface. Ligands are represented in cyan (docked pose in the presence of SRM), pink (docked pose in the absence of SRM) and yellow (crystal structure) ball and stick. In figure A, C and E the docked pose is perfectly aligned with the crystal ligands. SRMs are represented in grey surface model.

In the case of 2I4H, the active site features a phosphate-binding element formed by the backbone residues from 1904 to 1910 and the side chain of D1910[23]. In the crystal structure, the ligand engages in hydrogen bonding with key residues such as R1910, S1905, V1908, and A1906, which are crucial for stabilizing the ligand within the active site. Additional active site residues, including D1910, C1904, V1908, Q1948, and H1871, also significantly contribute towards the ligand interactions. The phosphate-binding site, flanked by residues N1735, Y1733, and I1736, plays a pivotal role in ligand binding, with the p-ethyl phenylsulfamic acid moiety of the ligand forming hydrogen bonds with H1871, D1870, and R1910. The H1871 is particularly notable for its π-stacking interaction with the ligand’s phenyl ring moiety. The enzyme’s WPD-loop undergoes a significant conformational change upon ligand binding, facilitating the proper alignment of the catalytic residue D1870 and ensuring precise ligand fit within the active site**(Figure 3A)** [24].

In contrast, in the docked structure without SRM, the p-ethyl phenylsulfamic acid adopts the same conformation as in the crystal structure due to being captured in a small cleft of the protein. However, the rest of the ligand orients entirely in the opposite way when compared to the crystal pose **(Figure 3B)**. Despite this, the docked structure maintains hydrogen bonding with several key residues such as S1905, A1906, V1908, G1909, R1910, D1870, and H1871. In the absence of SRM, the π-stacking interaction with H1871 is absent. This absence likely affects the proper alignment and conformational transition of the WPD-loop, in turn resulting in increased RMSD values for the ligand.

In the case of 5EI4, the WPF shelf—a conserved triad of tryptophan, proline, and phenylalanine—provides a hydrophobic platform that stabilizes ligand binding. The gatekeeper residue, isoleucine, shapes the acetyl-lysine binding pocket, influencing ligand access and contributing to selectivity and affinity. In the crystal structure, the ligand binds to the enzyme through several key interactions. The triazole moiety of the ligand forms a hydrogen bond with the residue D88, anchoring the ligand securely within the binding pocket. The triazolopyrimidinyl fragment is oriented within the ZA channel (formed by the Z and A helices in the protein) and forms van der Waals contacts with residues L92 and Q85, along with hydrogen bonds with N140. The pyrimidine ring in the ligand strongly interacts with Q85[25]. These interactions provide specificity for the ligand towards the active binding domain **(Figure 3C)**.

In the absence of SRM, the docked structure reveals an altered interaction pattern. The triazolopyrimidinyl fragment and triazole moiety are oriented in the opposite directions compared to the crystal structure. This orientation leads to a different pattern of hydrogen bonds between the ligand and active site residues. In the docked pose, different portions of the ligand form hydrogen bonds with N140 and D88. These interactions might be less effective, likely contributing to the higher RMSD, causing the ligand to adopt a conformation that deviates significantly from the optimal binding pose observed in the crystal structure **(Figure 3D)**.

In 1Q0Z, the active site is located at the interface of two domains. The ligand binds with its hydrophobic part extending deep into the pocket and its carbohydrate moiety at the entrance. Hydrophobic and stacking interactions occur between the ligand and the residue F134. The primary amino sugar forms additional interactions with residues M103, H219, Y220, and L222, and a hydrogen bond with D135 **(Figure 3E)** [26].

While analyzing the docked pose in the absence of SRM, the ligand exhibited similar interactions within the binding cavity of the proteins. Specifically, a hydrogen bond with D135 and a π-stacking interaction with F134 were noted, consistent with the interactions seen in the presence of SRMs. However, the absence of SRM allowed more conformational freedom to the sugar units protruding towards the solvent. This increased flexibility led to an altered conformational orientation of this portion of the ligand, as reflected in the higher RMSD values. This deviation indicates that the ligand adopts a different binding pose in the absence of SRM, which may impact its binding affinity and overall stability within the active site **(Figure 3F)**.

The absence of SRMs appears to impact the stability and accuracy of ligand binding, altering interaction patterns and increasing RMSD values. These findings suggest that SRMs play a critical role in stabilizing key interactions within the protein-ligand complex, which in turn enhance the docking protocols and improve the accuracy of ligand binding predictions in computational studies. However, in the specific cases of PDB IDs: 6G0G, 6GL8 and 5ACL, we observed only minimal deviations for the ligands even in the absence of SRMs **(Figure 4A to C)**. To delve deeper into this phenomenon, we conducted a comprehensive structural analysis to understand the reasons behind the reduced RMSD for these molecules. Our investigation revealed that these ligands are confined within a relatively shallow and linear binding pocket. This particular structural characteristic of the binding site significantly influences ligand behavior, where the ligands adopt a parallel orientation within these binding sites (parallel orientation is with respect to the binding site), which inherently restricts their movement of ligand within the binding pockets, resulting in lower RMSD values.

**Figure 4.**
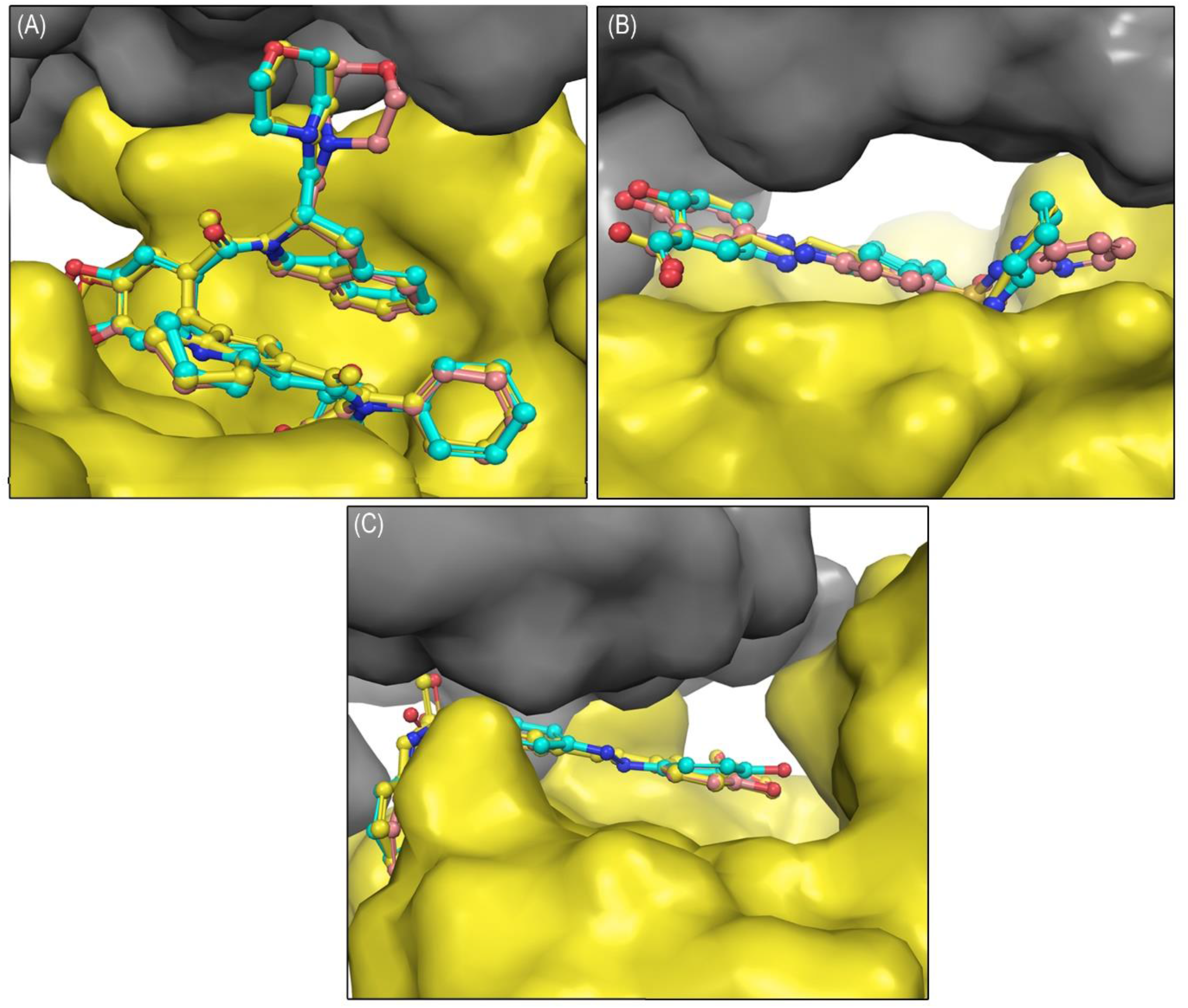
Structural comparison of the binding modes of the selected candid ates: Panels A-C depict the interactions of ligands within the PDB entries 6G0G, 6GL8, and 5ACL, respectively. In each case, the SRM is illustrated as a grey surface. The docked poses, both in the presence (cyan) and absence (pink) of SRMs, are superimposed onto the crystal structure, which is displayed in yellow. The ligands are represented using ball-and-stick models.

### MD Simulations confirmed the importance of SRMs

To gain deeper insights into the dynamic behaviour of protein-ligand complexes, MD simulations were performed under two distinct conditions: in the presence of SRMs **(case-1)** and in their absence **(case-2)**. For the simulations, three specific structures were selected based on their highest RMSD values observed after the docking step in the absence of SRMs: PDB IDs 2I4H (RMSD = 7.64Å), 5EI4 (RMSD = 6.63Å), and 1Q0Z (RMSD = 2.97Å). The rationale behind selecting these structures was to investigate if the ligands could achieve the crystal conformation by undergoing structural rearrangement during the MD simulations. Upon comparing the docked poses of case-1 and case-2 with the crystal pose, it was observed that case-2 exhibited a higher RMSD than case-1. This increased RMSD in case-2 is likely attributed to the enhanced conformational freedom of the ligand regions exposed to the solvent surface. In contrast, in case-1, the presence of SRMs significantly restricts this conformational freedom of the ligand that is exposed to the solvents, leading to a more stable ligand conformation.

To our surprise, none of the selected ligands achieved the crystal conformation during MD simulations. However, these ligands exhibited very low RMSD profiles throughout the simulations. During the 100 ns simulations, the stability of both the protein and the ligands was well maintained, even in case-2 **(Figure 5)**. The lower RMSD observed in case-2 systems can be attributed to the other stable interactions formed between the ligands and the protein over the simulation period. We also assessed the ligand dynamics for both case-1 and compare with that of the crystal poses, revealed very low RMSD profile. It should be noted that the binding pose selected for the MD simulations was determined through molecular docking studies. This suggests that SRMs play a crucial role in maintain the biologically relevant/crystallographically significant pose.

**Figure 5.**
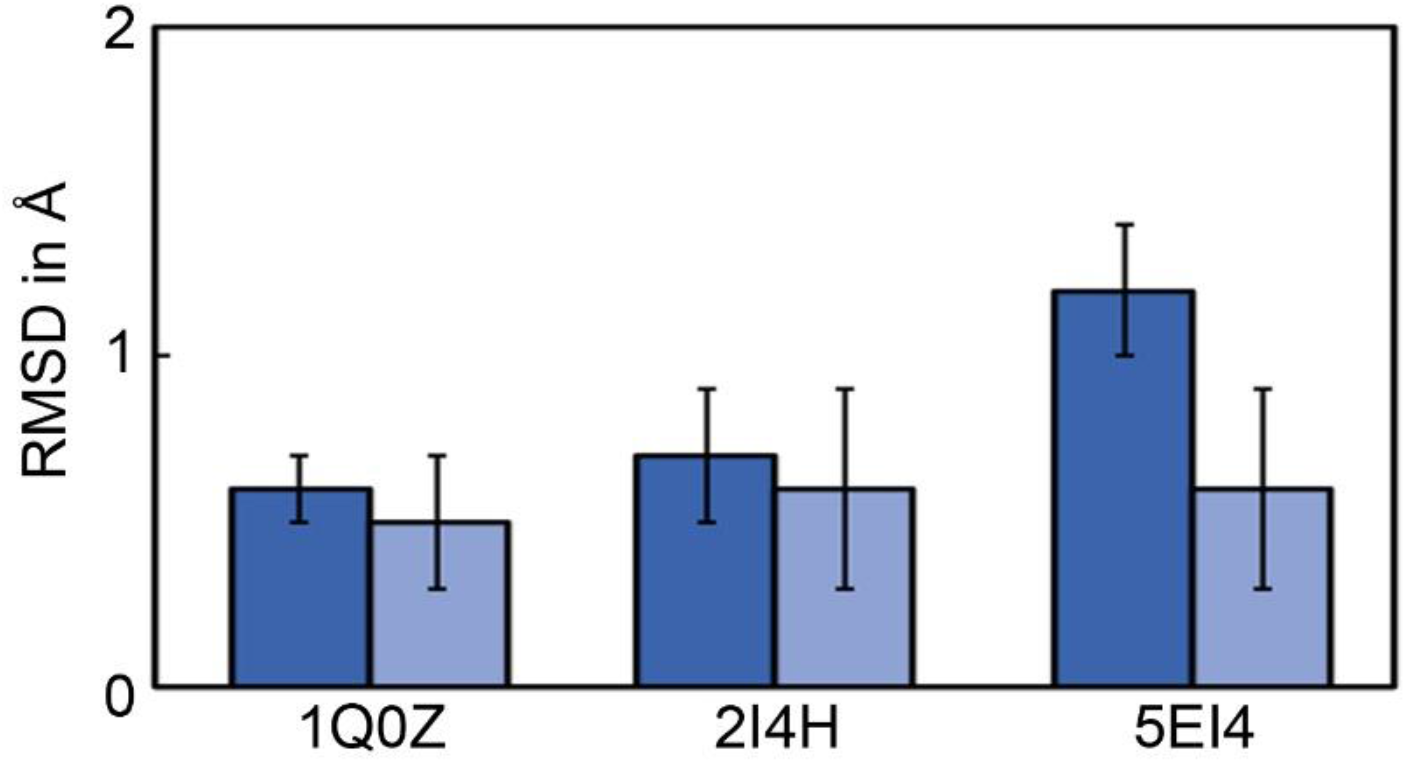
RMSD of ligands in the selected PDB entries such as 1Q0Z, 2I4H and 5EI4 during 100 ns simulations. In the figure, the dark blue represents the presence of SRMs, while light blue represents their absence.

A critical analysis of docked poses and MD trajectories was carried out to understand the differences in binding modes between case-1 and case-2. In the case of 1Q0Z, both case-1 and case-2 reproduced almost the same interactions between the ligand and the deeper cleft of the binding pocket. A common hydrogen bond with D135 and a face-to-face stacking interaction with F134 were observed in both case-1 and case-2 throughout the MD simulation, resulting in a stable orientation of the ligand inside the binding cavity. However, the differences between case-1 and case-2 became apparent in the protruding part of the ligand. In case-1, additional interactions were observed with residues from the SRM, such as A196, E197, and S280, which stabilized the ligand in a pose very similar to the crystallographic one. In contrast, in case-2, the formation of a hydrogen bond with L222 led to a different orientation of ligand, which remained stable throughout the MD simulation **(Figure 6A)**. This detailed comparison highlights how the presence of SRMs in case-1 reinforces the primary interactions within the binding pocket to adopt a conformation close to the crystallographic pose. Conversely, in the absence of SRMs, as seen in case-2, the ligand adopts a different stable orientation due to alternative interactions, emphasizing the significant role of SRMs in modulating ligand conformation and stability within the binding site **(Figure 6B)**.

**Figure 6.**
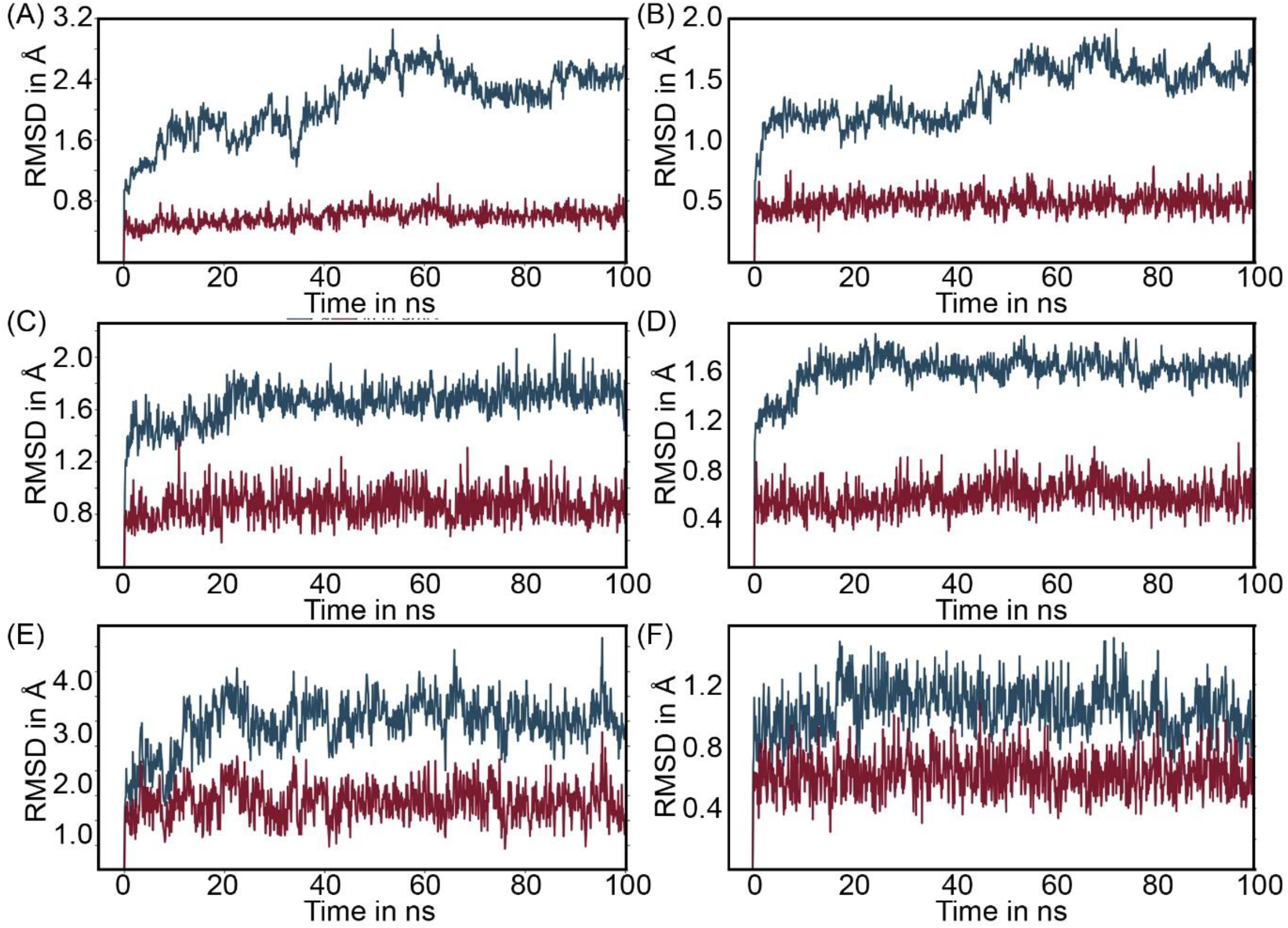
RMSD of selected proteins (all blue lines) and ligands (all red lines) over 100 ns simulations are shown. The docked poses of 1Q0Z, 2I4H, and 5EI4 in the presence (figures A, C, and E, respectively) and absence (figures B, D, and F, respectively) of SRM are displayed. In all figures, time is shown on the X-axis and RMSD on the Y-axis.

**Figure 7.**
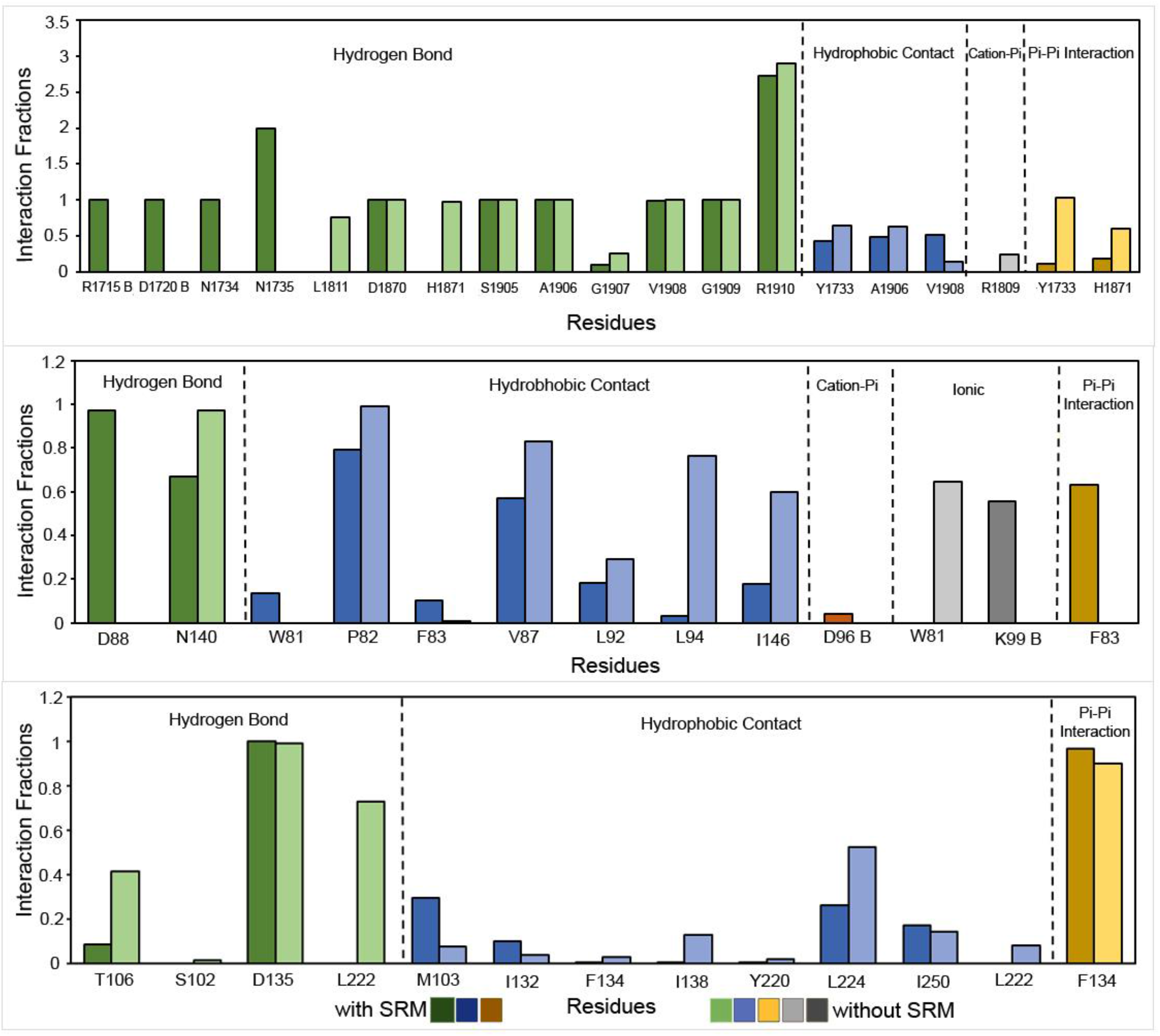
The intermolecular contacts formed between the ligand and proteins during simulation in the selected structures. (A) The RMSD of ligands during MD simulations with reference to. the crystal poses in the presence (dark blue) and absence (light blue) of SRMs. (B-D) Protein ligand interactions fractions in 2I4H, 5EI4 and 1Q0Z.

In the case of 5EI4, the simulations for case-1 accurately reproduced the binding pose of the ligand ‘5NV’ within the binding pocket. The ligand’s two moieties, 1H-1,2,4-triazol-3-amine (moiety-1) and chlorobenzene (moiety-2), were observed to protrude towards the solvent surface. In case-1, the orientation of moiety-1 was determined by a strong hydrogen bond with D88 and an interaction with D96 from the SRM, while moiety-2 was stabilized primarily by a π-π stacking interaction with K99 from the SRM. In contrast, in case-2, these two moieties underwent a ∼50° flip to the opposite side compared to the crystal pose resulting into a stacking interaction with residue W81 through moiety 1 and strong hydrogen bond with N140. This reorientation and the resulting stable interactions were further supported by the RMSD values extracted from the MD simulations of case-1 **(Figure 6C)** and case-2 **(Figure 6D)** relative to the crystal pose. In case-1, the presence of SRMs facilitated more extensive and stable interactions, closely mirroring the crystallographic pose. Conversely, in case-2, the absence of SRMs led to alternative interactions and a notable reorientation of the ligand, demonstrating the critical role of SRMs in maintaining ligand conformation and stability within the binding pocket.

In the case of 2I4H, the simulations of both case-1 and case-2 reproduced a stable hydrogen bond interaction between the ligand ‘UA1’ and residues located deep inside the active site, including R1910, G1909, V1908, A1906, S1905, and D1870. Similar to the observations in 1Q0Z, the protruding part of the ligand adopted a different orientation in case-2. In case-1, a stable hydrogen bond with N1734, along with interactions with R1715 and D1720, stabilized the ligand in a position closely resembling that of the crystallographic pose. These interactions effectively anchored the ligand within the binding site, minimizing conformational changes and maintaining a low RMSD. Conversely, in case-2, the ligand’s portion which is protruding towards the solvent formed hydrogen bonds with K1811 and H1871, and exhibited a classical π-π stacking interaction. These interactions resulted the docked pose in a different orientation when compared to the crystal pose. Despite this deviation, all interactions remained stable throughout the simulation period, which was reflected in the lower RMSD for this ligand. In case-1, the presence of SRMs not only reinforced primary interactions but also introduced additional stabilizing contacts that guided the ligand towards a conformation similar to the crystallographic pose **(Figure 6E)**. In contrast, case-2 demonstrated that in the absence of SRMs, the ligand achieved stability through alternative interactions, leading to a different but stable orientation. These findings highlight the dynamic interplay between ligands and their binding environments, influenced significantly by the presence or absence of SRMs **(Figure 6F)**.

## CONCLUSION

The experimental determination of ligand binding is crucial in the drug discovery process. Techniques such as X-ray crystallography, cryo-electron microscopy (cryo-EM), and nuclear magnetic resonance (NMR) spectroscopy are essential for elucidating the three-dimensional structures of protein-ligand complexes. These methods provide invaluable insights into the molecular interactions that underpin drug efficacy and specificity. However, accurately determining ligand binding modes presents significant challenges due to the limitations in achieving high-resolution structures and the presence of structural ensembles. Often, computational approaches are employed to overcome these hurdles and achieve a more precise understanding of ligand binding dynamics.

Molecular docking and dynamics are considered promising approaches for predicting ligand binding. However, predicting/reproducing the crystal pose is sometimes challenging in the computational drug discovery. The difficulty lies in accurately simulating how a ligand fits into a receptor’s binding site in a biologically significant manner. Despite advances in algorithms and computational power, achieving reliable pose predictions is still a complex task due to the dynamic nature of protein-ligand interactions and the vast conformational space that must be explored.

To improve the accuracy of these predictions, we propose a novel approach that involves incorporating symmetry-related molecules into the docking process. This strategy leverages the inherent symmetrical properties of certain molecular complexes to refine the docking results. Our hypothesis is that this method will enhance the precision of pose predictions, particularly for ligands with a solvent-accessible surface area (SASA) greater than 15% and that engage in direct interactions with symmetry-related molecules.

To test the efficacy of this approach, we conducted a series of validations using a small but diverse dataset. Our findings indicate that ligands arranged in parallel orientations within shallow binding pockets, which still maintain interactions with symmetry-related molecules, exhibit minimal impact on their conformational flexibility. This observation suggests that such ligands possess inherently rigid degrees of freedom, which reduces the variability in their binding conformations and enhances the robustness of the docking predictions.

Overall, our study underscores the potential of incorporating symmetry-related molecules in molecular docking to achieve more accurate and biologically relevant pose predictions. Further research and larger-scale validations are necessary to fully understand the implications and potential limitations of this approach, but our initial results are promising and warrant deeper investigation.

## ACKNOWLEDGEMENT

R.C. gratefully acknowledges the financial support received from Kerala State Higher Education Council, Govt. of Kerala in the form of Nava Kerala post-doctoral fellowship.

